# Leveraging a genetically tractable alphaproteobacterium reveals molecular determinants of bacterial growth in fungal-decayed wood

**DOI:** 10.64898/2026.05.07.723453

**Authors:** Nathan Lewis, Irshad Ul Haq, Jonathan S. Schilling, Kathryn R. Fixen

## Abstract

Brown rot wood-degrading fungi release carbon (C) from deadwood but leave behind a large fraction of C sequestered in lignin residues or as fungal metabolites. The strength of sequestration in these C residuals remains unclear, but proteobacteria-dominated bacterial communities have been implicated in metabolizing C from decay residues, possibly erasing the C sequestration potential assumed for brown rot. Here, we paired a model brown rot fungus (*Rhodonia placenta*) with a model Alphaproteobacterium (*Rhodopseudomonas palustris*) to track fungal release and bacterial utilization of C derived from decaying wood. We found that fungal decay products generated by *R. placenta* could be used by *R. palustris* for growth, and later decay stages contained more usable substrates than early stages. High performance liquid chromatography with mass spectrometry identified a range of aromatic and non-aromatic compounds in the fungal-decayed wood, but after 95 days of bacterial growth, *R. palustris* preferentially consumed non-aromatic acids over aromatic lignin monomers. Genes involved with aromatic compound degradation were unimportant for bacterial growth, and RNA sequencing revealed that aromatic compound degradation genes were repressed on decayed wood extract. Randomly barcoded transposon sequencing failed to identify a solitary catabolic pathway used by *R. palustris*, suggestive of substrate co-utilization, and surprisingly, showed that genes involved with copper toxicity were essential. Finally, we found that genes involved with biosynthesis of certain cofactors and amino acids were no longer essential on decayed wood extract, suggesting these nutrients were readily accessible. This study helps lay the foundation to understand potential bacterial-fungal interactions in decayed wood.

**Graphical abstract:** To explore how brown rot fungi support and compete with bacterial partners in the wood decay environment, the model brown rot fungus *Rhodonia placenta* was used to degrade aspen wafers which were then infused into bacterial growth medium. By leveraging the range of molecular biology tools available for the model Alphaproteobacterium *Rhodopseudomonas palustris*, we discovered that *R. palustris* preferentially consumes short organic acids instead of aromatic lignin monomers which it would otherwise consume if provided in isolation. Additionally, *R. palustris* scavenged certain amino acids (AAs) and enzyme cofactors including methionine, biotin, and PLP from the decayed wood extract, highlighting these as key shared resources for bacterial-fungal partnerships. We found that *R. placenta* increased the concentration of certain metals (Cu and Al) inducing a metal stress response in *R. palustris*, indicating that metal toxicity could be an important mode of competition between fungi and bacteria in the wood decay environment.

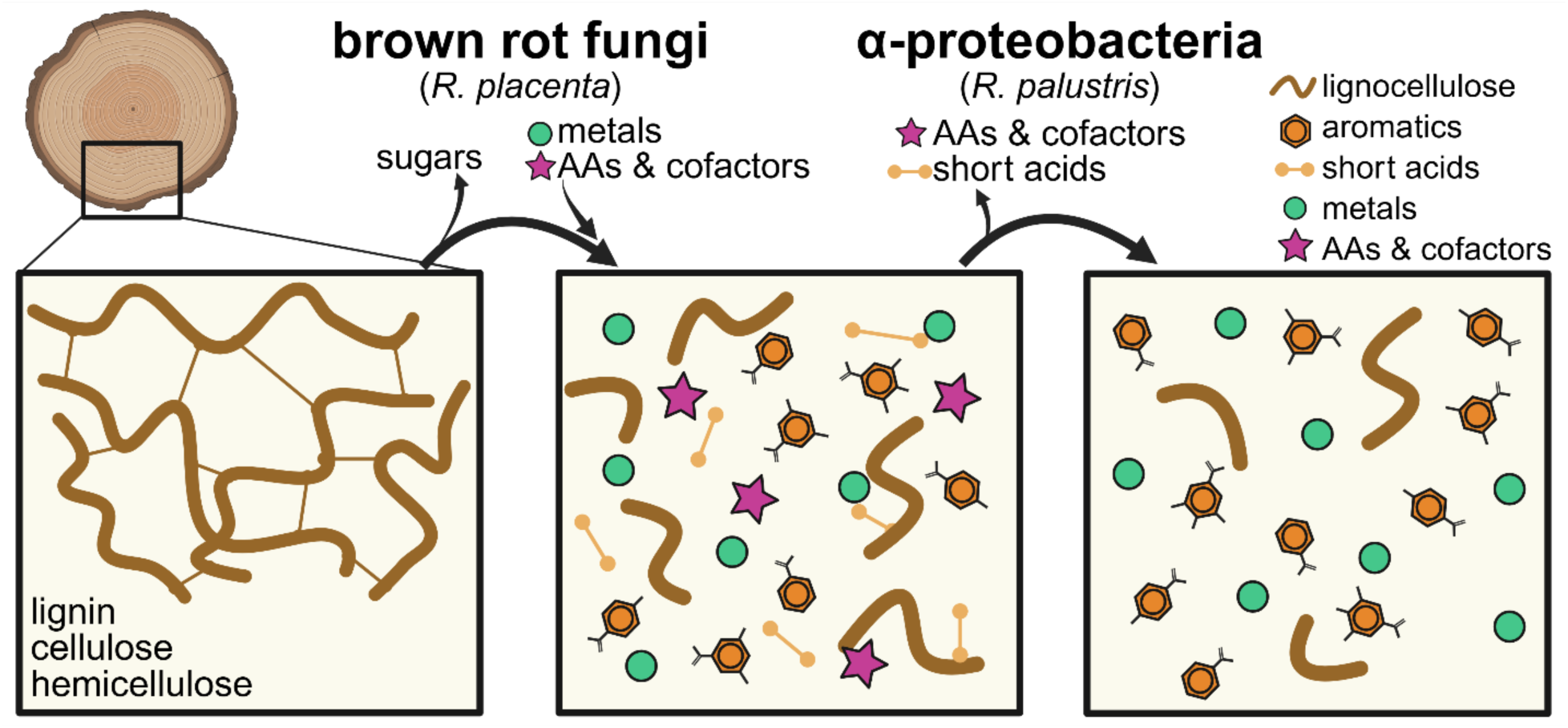

## Introduction

Wood rot fungi decompose lignocellulose and play an important role in carbon cycling in natural environments [1]. Wood rot fungi also help shape the microbial communities found in decomposing wood by altering the availability of metabolites, and distinct decay mechanisms have been shown to affect bacterial community composition [2]. Interactions between wood rot fungi and bacteria in these environments is not well understood, but there are indications of both antagonistic and beneficial interactions in decomposing wood communities [3]. Understanding these interactions is of ecological and economic importance, as they likely play a role in controlling the rate and the fate of carbon released from a massive global pool of biotic carbon. Addressing these interactions requires approaches that directly interrogate the genetic and metabolic determinants of microbial interactions in wood decay environments.

High-throughput sequencing approaches such as transcriptomic sequencing (RNA-seq) and transposon sequencing (Tn-seq) have transformed our ability to study microbial interactions at the molecular level [4]. Such approaches can validate predicted interactions and uncover pathways involved in these interactions that may not be predicted *a priori* [5–7]. Tn-Seq is particularly effective for identifying potential cross-feeding interactions in microbial communities, providing insight into the genetic basis of cooperative and competitive interactions [6,8,9]. When community members are not genetically tractable, Tn-seq libraries of model organisms can be used to probe interaction-relevant environmental features. This strategy has been successfully applied in cheese rind biofilm communities, where the model bacterium, *Escherichia coli,* was used to interrogate interactions with other community members, including fungi [5].

A similar strategy can be applied to investigate how fungal wood decay processes shape local bacterial communities. Here, we focused on the model brown rot fungus *Rhodonia placenta* together with the phototrophic Alphaproteobacterium *Rhodopseudomonas palustris*. *R. placenta* is one of the most well-studied wood-degrading fungi and is an ideal system to understand fungal-bacterial interactions. Unlike white rot fungi that use Class II peroxidases to target and degrade lignin, brown rot fungi like *R. placenta* use extracellular Fenton chemistry to create hydroxyl radicals, which depolymerize cellulose and hemicellulose and leave lignin modified but largely intact [10,11]. The degradation profile at different decay stages has been recently characterized using metabolomics for *R. placenta* and includes the release of organic acids and aromatic compounds that could be consumed by bacterial partners [12].

Although traditional wood decay community studies have largely focused on fungi, the studies that have included a co-assessment of bacteria share a common theme of Alphaproteobacterial dominance, and occasionally identify certain *Rhodopseudomonas* species [2]. *R. palustris* is a phototrophic facultative anaerobe capable of metabolizing a wide range of organic acids and aromatic compounds, including lignin derivatives, and has been shown to metabolize these compounds in corn stover hydrolysate [13]. *R. palustris* is also a well-studied model organism with a suite of reliable genetic tools which can be used to probe nutrient availability and potential interaction-relevant features of the decay environment. RNA-seq and Tn-seq have been successfully applied to *R. palustris* to detect changes in gene expression and gene fitness across diverse environmental conditions [14,15]. The combination of metabolic versatility and genetic tractability makes *R. palustris* an ideal model bacterium to investigate potential fungal-bacterial interactions in decomposing wood.

In this study, we found that *R. palustris* can grow using substrates from early- and late-stage fungal decay of *R. placenta* as the sole carbon source. Although aromatic compounds were detected in the decayed wood, genes required for aromatic compound degradation were not necessary for *R. palustris* to grow in decayed wood extract and expression of most genes involved in aromatic compound degradation were not induced by decayed wood extract. Instead, *R. palustris* consumed various non-aromatic organic acids. Randomly barcoded transposon sequencing (RB-TnSeq) revealed that *R. palustris* tolerated insertions in vitamin and amino acid biosynthesis genes when grown with decayed wood extract but not with acetate as a carbon source, indicating that these nutrients were acquired from the decayed wood extract. RB-TnSeq and RNA-seq also revealed a role for metal homeostasis in survival of *R. palustris* in wood decay extracts. Together, these findings provide insight into how metabolites or resources generated by brown rot fungi colonizing wood substrates can shape bacterial wood decay communities.

## Materials and Methods

### Growth of bacterial cultures

Cultures of *R. palustris* strains were grown in a phosphate buffered mineral medium referred to as photosynthetic medium (PM) whose formulation lacks carbon and energy sources so it can be supplemented with specific carbon sources as described below. The exact constitution of PM and general cultivation methodology has been described in other work [16,17]. All bacterial cultures, unless otherwise stated, were grown at 30°C in septum sealed Hungate-style tubes (Chemglass life sciences, Vineland, NJ) with an anaerobic atmosphere (97.5% N_2_ and 2.5% H_2_) within 5.5 inches of a 60 W incandescent light bulb supplying 30 μmol photons m^−2^s^−1^ as an energy source [16,18]. Starting cultures of *R. palustris* CGA009 were inoculated into 10 mL of liquid PM supplemented with 20 mM acetate as a carbon source and 0.1% yeast extract to supply additional vitamins. After reaching stationary phase (OD_660_ = 1.1) 500 μL of the starting culture was removed and transferred to a 1.5 mL tube, then centrifuged (2 minutes at 10,000 g and 4 °C) to pellet the cells. Unless otherwise specified, all centrifugation was performed using these parameters. Pelleted cells were resuspended with PM lacking any carbon source or yeast extract, then centrifuged and resuspended with PM again to remove any residual carbon substrates from the starting culture medium. Washed cells were used to inoculate 10 mL of anoxic liquid PM supplemented with 20 mM acetate or supplemented with decayed wood extract (described below) as a carbon source. These cultures were inoculated to an initial OD_660_ of 0.03 and incubated in the light.

Because the exact concentration of biologically available carbon in the decayed wood extracts is difficult to determine, several preliminary experiments were performed to identify the optimal volume of wood extract to add to bacterial cultures. In trials, we found that adding 2 mL of wood extract generated final cell densities slightly lower than a culture containing 20 mM acetate. To prepare cultures with wood extract as a carbon source, 2 mL of medium was removed from sealed 10 mL Hungate tubes, then 2 mL of wood extract was added to bring the final volume back to 10 mL. The solvent for the wood extraction procedure was sterile PM to prevent dilution of medium components. The wood extraction procedure is described below in greater detail.

### Fungal strains, media, and soil-block microcosms

*Rhodonia placenta* strain MAD 698-R (American Type Culture Collection 44394) was maintained on malt extract agar for routine culturing. For establishment of wood decomposition microcosms, the American Wood Protection Association (AWPA) standard E10-16 (2016) was used with modification. Briefly, we used a 1:1:1 mixture of fertilizer-free soil, peat, and vermiculite, which was hydrated to a moisture content of 35-40% (wt/vol). The resulting hydrated mixture was packed into pint glass jars to one-third full. In each jar, two birch feeder strips (40 mm × 10 mm × 2 mm) were placed on the surface of the soil. Jars were covered with foil and autoclaved twice for 1 h (121 °C, 103 kPa) with the second cycle starting 24 hours after the first. After allowing the jars to cool for 24 hours, jars were inoculated with *R. placenta* by placing 1 cm agar plugs from malt extract agar plates into the jars. Soil jars were incubated aerobically at 26 °C for two weeks to achieve a confluent mycelial mat on the surface of the soil.

Aspen wafers (60 mm x 25 mm x 2.5 mm) were then placed flat on top of the *R. placenta* mycelial lawn and incubated aerobically at 26 °C with relative humidity maintained at 65 to 75%. Aspen (*Populus tremuloides*) was used for these experiments because it is an angiosperm with predictable and well-characterized lignocellulose chemistry consisting of (49% glucan, 17% xylan, 2.1% mannan, 0.5% arabinan, 2% galactan, 21% lignin, 3.7% acetyl, 4.3% uronic acids, 0.4% ash [19,20], and 3-4% total “extractives” [21]. For undecayed controls, soil-block jars including aspen wood wafers were prepared and incubated under the same conditions but were not inoculated with *R. placenta*.

### Preparation of decayed wood extracts

Decayed aspen wafers were aseptically transferred to 50 mL conical tubes and crushed using sterilized forceps to a fine powder. Undecayed wood retained its structural integrity and could not be crushed easily, so the undecayed wood was made into a powder by filing the wood to create a fine powder. The mass of wood powder was measured, and PM was added to the wood powder at a ratio of 10 mL of PM per gram of wood powder. The suspension was vortexed for 1 minute to thoroughly mix, then incubated overnight at 4°C to allow any soluble compounds from the wood to dissolve into the medium. The suspension was centrifuged for 30 minutes at 4000 x g and 4°C to separate the supernatant from the wood powder. The supernatant was poured onto a vacuum driven 0.22 μm PES membrane filter (Genesee scientific) and filter sterilized. The filter-sterilized wood extract was then transferred to a sterile 50 mL serum vial and sealed with a rubber stopper and aluminum crimp seal, then sparged with nitrogen for 30 minutes to remove dissolved oxygen from the wood extract. The filter-sterilized, oxygen-free wood extracts were stored at 4°C until used for culturing *R. palustris* as described above.

### Data availability

Transcriptomic data is available from the gene expression omnibus (GEO) under accession number GSE311457. Gene fitness data is available in the Genome Fitness browser (https://fit.genomics.lbl.gov/cgi-bin/myFrontPage.cgi) [22].

### Additional methods

Additional methods are supplied as supplemental material.

## Results

### Wood decayed by R. placenta generated carbon substrates consumed by R. palustris

Aspen wafers decayed by *R. placenta* to an early decay stage (28% mass loss, ‘Sollins’ decay class #2), a late decay stage (76% mass loss, decay class #5) [23,24] or not inoculated with *R. placenta* (undecayed) were infused into bacterial growth medium as described in the methods. Acid-insoluble (Klason) lignin analysis of late decay samples showed that the average mass loss was 76 +/− 2% (n = 7) and the average lignin loss was 44.6% +/− 0.1% (n = 7), giving a ratio of lignin loss to total mass loss of 0.58 (+/− 0.05, n = 7) which is characteristic of brown rot-dominated decay [25–27]. These early decay, late decay, and undecayed wood extracts were supplied to *R. palustris* in place of a carbon source in bacterial growth medium and used to grow *R. palustris* anaerobically with light as an energy source. *R. palustris* grown with early or late decay stage wood extract reached higher cell densities than samples provided with undecayed wood extract, and *R. palustris* reached the highest cell density with extracts from the late decay stage (Figure 1). This indicates that fungal activity was required to generate carbon substrates for growth of *R. palustris* and that more substrates are released as decay progresses.

**Figure 1:**
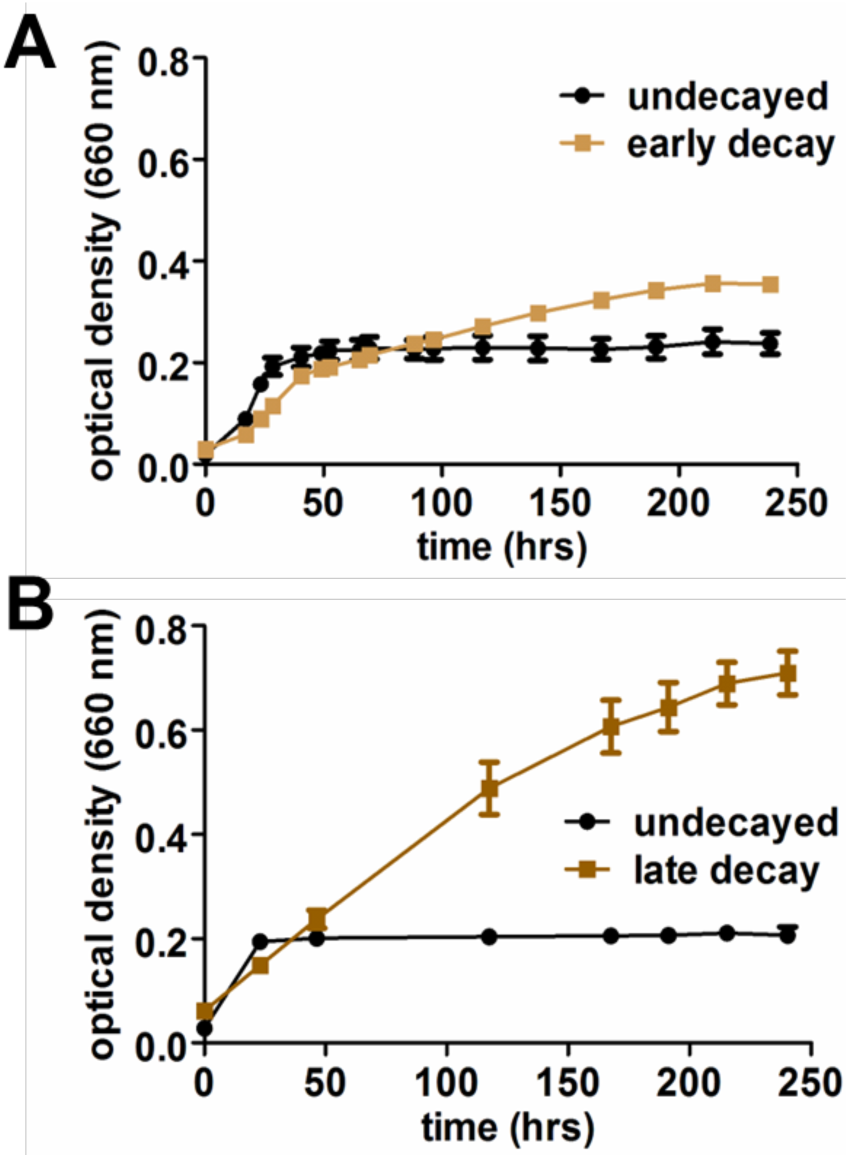
Brown rot fungal decay by *R. placenta* generates growth substrates for *R. palustris*. **A**. Growth curve of *R. palustris* CGA009 with undecayed wood extract (black) and early decayed wood extract (khaki). **B**. Growth curve of *R. palustris* CGA009 cultured with undecayed wood extract (black) and late decayed wood extract (brown). For both **A** and **B**, cultures were grown in anaerobic phototrophic conditions in biological triplicate, points represent the mean optical density for all three replicates and error bars represent one standard deviation above and below the mean.

Figure 1 also reveals that *R. palustris* does not grow exponentially on the decayed wood extracts, suggesting it is co-utilizing multiple substrates. To help identify what substrates were present in the late decay extract before and after the growth of *R. palustris*, we employed liquid chromatography coupled to mass spectrometry (LC-MS). *R. palustris* can grow with a wide variety of organic acids and aromatic compounds, including *p*-hydroxyphenyl (H), guaiacyl (G), and syringyl (S) types of aromatics, but it does not consume sugars like glucose, starch, or xylose [13,14,28–33]. Using LC-MS, we found that the decayed wood extract contains a range of organic acids and aromatic compounds (Table 1). Many of the compounds detected by LC–MS, such as benzoic acid, hydroxybenzoic acid, and coumaric acid, are assimilated by *R. palustris* when provided in isolation in similar growth conditions [29,31,32,34]. Although a variety of S-, H-, and G-type lignin monomers were identified, several aromatic compounds including benzoic acid, hydroxybenzoic acid, cinnamic acid, and vanillic acid were detectable even after 95 days of incubation with *R. palustris* (Table 1). In contrast, coumaric acid and the S-group lignin monomers sinapic acid and syringic acid were no longer detectable following bacterial growth. *R. palustris* cannot use syringic acid as a growth substrate [30,33] suggesting these compounds may undergo partial transformation rather than full assimilation, and compounds like sinapyl alcohol and sinapaldehyde were only detected after growth of *R. palustris.* Our LC-MS data shows that coumaric acid is absent after bacterial growth, and coumaric acid can be assimilated by *R. palustris* via the benzoate degradation pathway, so we hypothesized that the benzoate degradation pathway may be important for growth on the decayed wood extract.

**Table 1:** Presence or absence of specific compounds found in extracts of fungal decomposed wood before and after being used as a substrate for anoxic, photoheterotrophic bacterial growth.

|  | Compound | Decayed wood extract before growth <sup>a</sup> | Decayed wood extract after growth <sup>b</sup> |
| --- | --- | --- | --- |
| Short-chain acids | succinic acid | + | + |
|  | acetic acid | + | - |
|  | fumaric acid | + | + |
|  | oxaloacetic acid | + | - |
|  | oxalic acid | - | - |
| aromatic compounds <sup>c</sup> | hydroxybenzoic acid (H) | + | + |
|  | benzoic acid (B) | + | + |
|  | cinnamic acid (B) | + | + |
|  | vanillic acid (G) | + | + |
|  | coumaric acid (H) | + | - |
|  | syringic acid (S) | + | - |
|  | sinapic acid (S) | + | - |
|  | sinapyl amide (S) | + | + |
|  | sinapyl alcohol (S) | - | + |
|  | sinapaldehyde (S) | - | + |
|  | aminobenzoic acid | - | + |
|  | di-hydroxybenzoic acid (H) | - | - |

<sup>a</sup> ‘Before’ refers to the late-stage decayed wood extract in phosphate buffered mineral medium (PM medium) before *R. palustris* had been inoculated into the medium.
<sup>b</sup> ‘After’ refers to the cell-free spent culture medium after 95 days of anaerobic, phototrophic growth with *R. palustris*. Samples were taken from the growth curve shown in Figure 2 after the final measurement was recorded. As described in the methods, all cultures were maintained in septum-sealed anaerobic culture tubes until they were harvested for LC-MS analysis.
<sup>c</sup> The class of lignin monomer for each aromatic compound is identified using a letter in parenthesis following the name, where (H) indicates that the compound is a “Hydroxy” group lignin monomer, (G) indicates that it is a “Guiacyl” group monomer, and (S) indicates that it is a “Syringyl”
group monomer. The designation (B) is used for “Benzoic” to indicate that there are no aromatic ring substitutions aside from the C1 substitution.

To test this hypothesis, a strain of *R. palustris* which cannot assimilate aromatic compounds, including coumaric acid, was grown in minimal medium with late decay-stage wood extract as the carbon source. This strain has an in-frame deletion in *badE*, which encodes a structural component of benzoyl-CoA reductase, the central step in aromatic compound degradation [35]. *R. palustris* Δ*badE* grew similar to wild-type *R. palustris* when decayed wood extract was provided as a carbon source (Figure 2). The final optical density of the Δ*badE* mutant was only 10% lower than wild-type *R. palustris*, suggesting that it uses a majority of the same carbon source(s) as the wild type, and that aromatic compound degradation only explains 10% of the growth observed at maximum. These results indicate that, although aromatic compounds are present and may be transformed in the decayed wood extract, *R. palustris* does not primarily rely on their degradation for growth.

**Figure 2.**
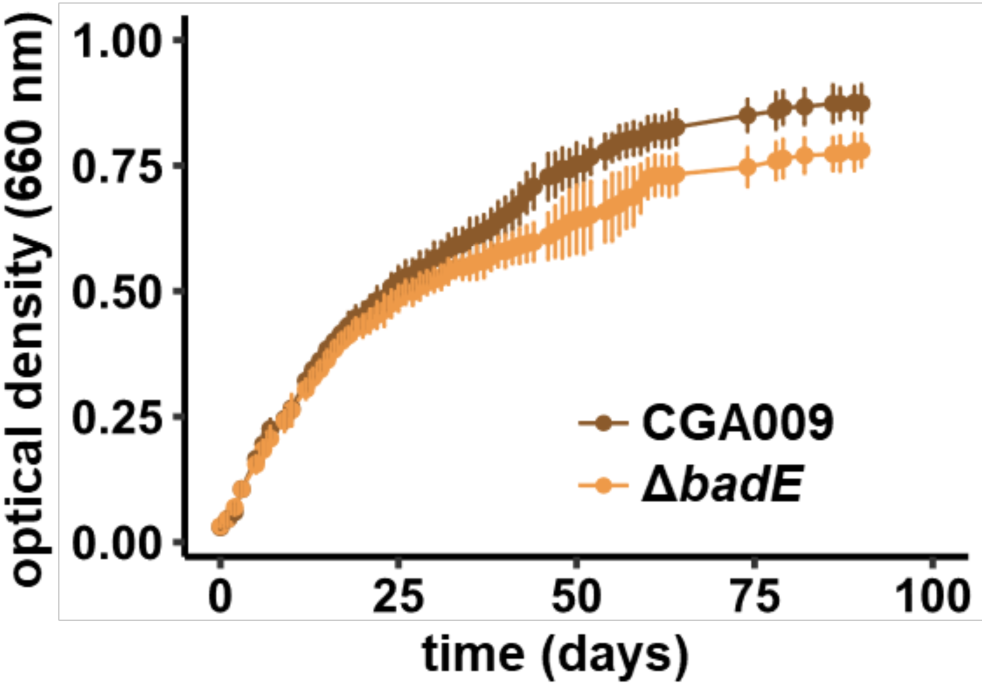
Benzoyl-CoA reductase is not necessary for growth on late decay wood extract. Both *R. palustris* CGA009 and *R. palustris* with an in-frame deletion in *badE* (Δ*badE*) were grown in minimal medium supplemented with late decay wood extract in anoxic phototrophic conditions. Each point represents the average of three biological replicates, error bars represent one standard deviation from the mean optical density.

To help explore what catabolic pathways or functions were being used for growth in the decayed wood extract, mRNA transcripts were collected from *R. palustris* grown in minimal medium with late decay wood extract as the carbon substrate and the same strain grown in minimal medium with acetate as a control. Transcriptomic sequencing (RNA-seq) was performed to determine which genes are differentially expressed in cells grown in the presence of the decayed wood extract compared to cells grown with acetate as a sole carbon source. Expression of most genes involved in aromatic compound degradation were not significantly differentially expressed (Supplemental dataset 1, Table 2). Only *vanB*, which encodes vanillate O-demethylase oxidoreductase, and *boxB*, a benzoyl-CoA oxygenase, were upregulated while the regulator *badR*, and two of the genes it regulates, *badI* and *badH*, were down-regulated (Table 2). Together, these data indicate that the limited induction of aromatic compound degradation pathways may contribute to the lack of detectable metabolism of aromatic compounds and supports the hypothesis that *R. palustris* primarily consumes other, non-aromatic carbon sources present in the decayed wood extract.

**Table 2.** Genes involved in aromatic compound degradation which are differentially expressed when decayed wood extract is provided as a carbon substrate.

| Locus tag<br>(RPA number) | Gene<br>name | Log<sub>2</sub> FC expression<br>(DE vs. AC)<sup>a</sup> | Annotation |
| --- | --- | --- | --- |
| TX73_018765<br>(RPA3621) | *vanB* | 4.3 | PDR/VanB family oxidoreductase |
| TX73_003505<br>(RPA0677) | *boxB* | 4.2 | benzoyl-CoA 2,3-epoxidase<br>subunit |
| TX73_003395<br>(RPA0655) | *badR* | -5.0 | MarR family transcriptional<br>regulator |
| TX73_003385<br>(RPA0653) | *badI* | -5.5 | 2-ketocyclohexanecarboxyl-CoA<br>hydrolase |
| TX73_003390<br>(RPA0654) | *badH* | -7.8 | 2-hydroxycyclohexanecarboxyl-CoA dehydrogenase |

<sup>a</sup>Positive values represent genes that are more highly expressed in decayed wood extract (DE) compared to acetate (AC). Negative values represent genes that are down-regulated in DE versus AC.

Brown rot fungi secrete organic acids such as oxalic acid, likely to acidify their microenvironment and promote Fenton chemistry [36–38]. In addition, lignocellulose contains ∼5% acetyl esters, releasing acetate during breakdown [39]. Consistent with this, several organic acid anions in the decayed wood extract were detected in LC-MS analysis, including succinate, acetate, and fumarate (Table 1). *R. palustris* is unable to metabolize sugars like glucose, sucrose, and xylose and grows best when consuming these short chain organic acids during photoheterotrophic growth [13,14,28]. After growth of *R. palustris*, acetate and oxaloacetate were no longer detected (Table 1). Therefore, *R. palustris* is most likely assimilating these short acids from the decayed wood extracts, preferring these substrates over the more recalcitrant aromatic compounds.

### RB-TnSeq suggests cofactors, vitamins, and oxidized carbon sources are accessible by bacteria

To gain insight into the genes essential for growth in these extracts and identify fungal resources which *R. placenta* could leave behind for its bacterial associates, a barcoded transposon library generated in *R. palustris* CGA009 was grown phototrophically in minimal medium supplemented with decayed wood extract as a carbon source. The library was also grown in minimal medium with 20 mM acetate as a carbon source so that genes required specifically for phototrophic growth with decayed wood extract could be differentiated from those required for phototrophic growth on acetate alone.

Comparing the fitness scores for each gene in both conditions revealed genes which are essential for growth with acetate but are no longer essential for growth when decayed wood extract is used as the carbon source (Figure 3). Transposon insertions in genes involved in synthesis of glycerol-3-phosphate (RPA0254), pyridoxine 5’ phosphate (RPA2694 and RPA3065), biotin (RPA2045, RPA2971, and RPA2970), methionine (RPA3702), and trehalose (RPA4661) were essential when acetate was supplied as a carbon source but were no longer essential when decayed wood extract was provided (Figure 3, Table 4) suggesting that these resources are being scavenged from the fungal-decay products.

**Figure 3.**
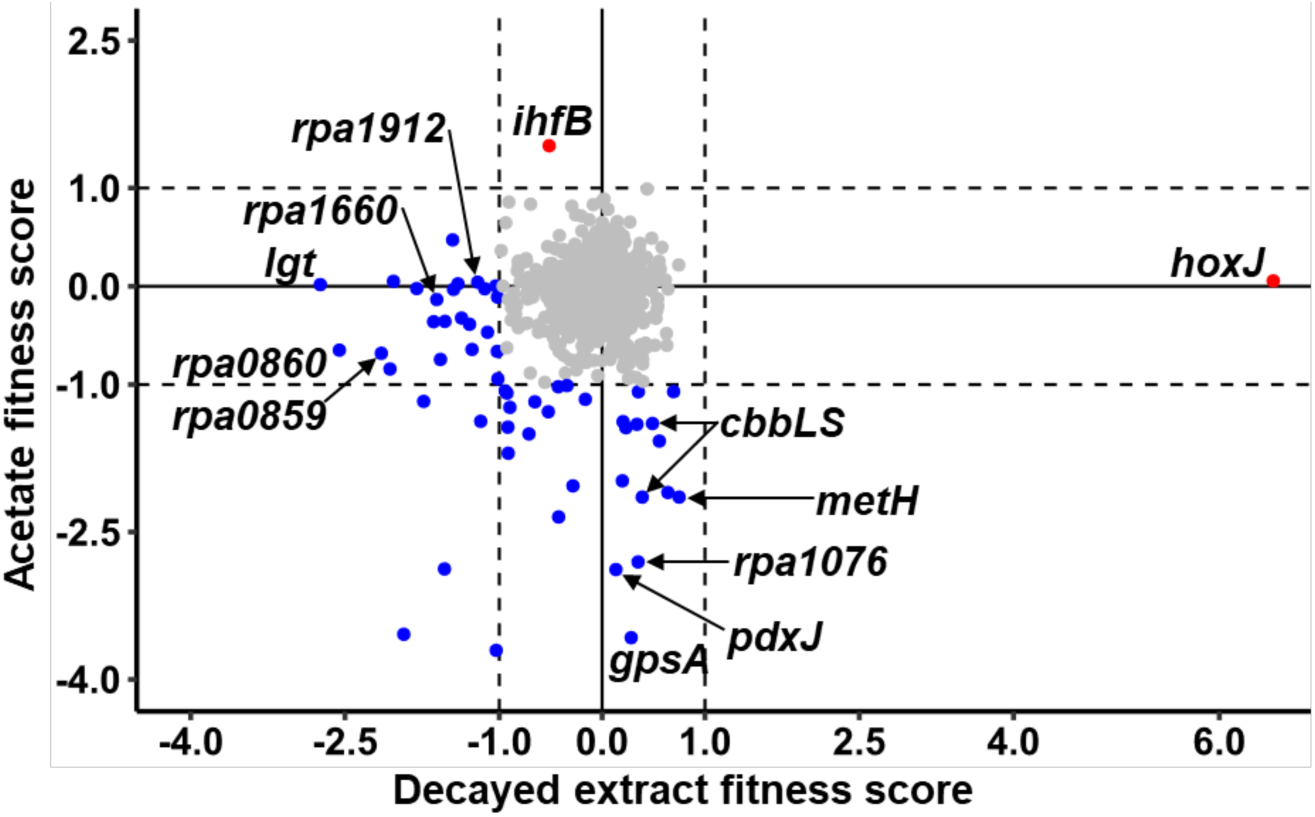
Comparison of fitness scores for *R. palustris* genes in minimal medium with acetate (Acetate fitness score) versus minimal medium with late decay-stage wood extract (Decayed extract fitness score). Each point represents the average calculated fitness score for three replicates. Relevant genes are labeled with arrows. Blue coloring indicates that a gene has a fitness score below −1 in either condition (fitness loss), red indicates that the gene has a fitness score above 1 in either condition (fitness gain), and gray indicates that a gene has a fitness score above −1 but below 1 (fitness neutral). Dotted lines represent a significance threshold of +/− 1 for either condition.

Transposon insertions in genes involved in redox homeostasis also showed improved fitness when grown with decayed wood extract compared to acetate. Carbon dioxide (CO_2_) fixation plays an important role in maintaining redox balance in *R. palustris* under photoheterotrophic conditions, particularly when growing with electron-rich carbon sources [40]. Transposon insertions in genes encoding the small and large subunit of type I RuBisCo (*cbbLS*) and a RuBisCo activase (*cbbX*) had a significant fitness defect when acetate was provided as a carbon source but did not have a significant fitness defect when decayed wood extract was provided (Figure 3). Additionally, insertions in *hoxJ* improved fitness of *R. palustris* in decayed wood extract but had no fitness benefit when acetate was provided. A strain with a deletion in *hoxJ* was previously shown to bypass a regulatory mutation that prevents expression of an uptake hydrogenase in *R. palustris* CGA009, enabling *R. palustris* to use hydrogen gas as an electron donor [41]. Together, these data indicate that *R. palustris* does not need to reduce CO_2_ to maintain redox balance and may benefit from an additional electron source like hydrogen when growing in decayed wood extract. This suggests that the decay extracts contain more oxidized carbon substrates that can be metabolized by *R. palustris*. Consistent with this, the oxidized carbon substrate fumarate was detected in the decayed wood extract (Table 1), and *R. palustris* is known to utilize fumarate as a carbon source [42].

### Genes involved in horizontal gene transfer and metal homeostasis are induced and required for growth in decayed wood extract

We next sought to identify genes whose disruption resulted in fitness defects during growth in late decay wood extract, thereby revealing pathways that play a role in survival under these conditions. Unexpectedly, *R. palustris* genes encoding a gene transfer agent (GTA) appear to play a role in late decay extracts. GTAs are phage-like particles that package random fragments of host genomic DNA and are released through cell lysis, facilitating horizontal gene transfer to recipient cells (reviewed in [43]). Increased expression of the genes required to assemble the GTA was observed in *R. palustris* grown in late decay extract (Table 3). RB-TnSeq analysis revealed that insertions in RPA1912, which encodes the baseplate component of the GTA tail structure, resulted in reduced fitness during growth on late decay extracts, suggesting that incomplete GTA assembly negatively impacts fitness (Figure 3 and Supplemental Dataset 2). These results provide the first evidence of GTA induction in *R. palustris* and suggest that horizontal gene transfer could be occurring and be important for growth under these conditions.

**Table 3.** Selected genes upregulated in *R. palustris* when decayed wood extract is provided as a carbon substrate.

| Locus tag<br>(RPA number) | Gene name | Log<sub>2</sub> FC gene<br>expression<br>(DW vs. AC)<sup>a</sup> | Annotation |
| --- | --- | --- | --- |
| *Copper response*<sup>b</sup> |  |  |  |
| TX73_004475 |  | 4.2 | hypothetical protein |
| TX73_004485<br>(RPA0870) |  | 5.6 | type III PLP-dependent enzyme |
| TX73_008540<br>(RPA1661) | *csoR* | 4.1 | metal-sensitive transcriptional<br>regulator |
| TX73_009940<br>(RPA1929) |  | 7.4 | Do family serine endopeptidase |
| TX73_009945<br>(RPA1930) | *cusR* | 4.0 | response regulator transcription<br>factor |
| TX73_010565<br>(RPA2050) |  | 5.4 | Spy/CpxP family protein refolding<br>chaperone |
| TX73_015000<br>(RPA2894) |  | 7.9 | hypothetical protein |
| TX73_018485 |  | 4.1 | hypothetical protein |
| TX73_023750<br>(RPA4572) | *htrA/degQ/degS* | 7.5 | Do family serine endopeptidase |
| *Denitrification* |  |  |  |
| TX73_010630<br>(RPA2062) | *nosD* | 4.1 | nitrous oxide reductase family<br>maturation protein NosD |
| TX73_010650<br>(RPA2066) | *nosX* | 3.5 | FAD:protein FMN transferase |
| TX73_019220<br>(RPA3710) | *nirA* | 9.2 | NirA family protein |
| *Gene transfer agent* |  |  |  |
| TX73_009710<br>(RPA1885) |  | 4.4 | phage portal protein |
| TX73_009715<br>(RPA1886) |  | 6.5 | hypothetical protein |
| TX73_009725<br>(RPA1887) |  | 3.7 | HK97 family phage prohead<br>protease |
| TX73_009730<br>(RPA1888) |  | 3.6 | phage major capsid protein |
| TX73_009745<br>(RPA1891) |  | 3.6 | hypothetical protein |
| TX73_009750<br>(RPA1892) |  | 4.3 | DNA-packaging protein |
| TX73_009770<br>(RPA1896) |  | 3.8 | head-tail connector protein |
| TX73_009775<br>(RPA1897) |  | 4.3 | head-tail adaptor protein |
| TX73_009780<br>(RPA1898) |  | 4.2 | DUF3168 domain-containing<br>protein |
| TX73_009785<br>(RPA1899) |  | 4.2 | phage major tail protein, TP901-1 family |
| TX73_009800<br>(RPA1902) |  | 4.0 | phage tail tape measure protein |
| TX73_009815<br>(RPA1905) |  | 3.9 | DUF2163 domain-containing protein |
| TX73_009820<br>(RPA1906) |  | 4.0 | C40 family peptidase |
| TX73_009850<br>(RPA1912) |  | 3.8 | baseplate multidomain protein megatron |
| TX73_009855<br>(RPA1913) |  | 4.5 | DUF2793 domain-containing protein |

<sup>a</sup> Log<sub>2</sub> of the fold change (FC) in normalized transcript counts for specific genes. Positive values represent genes that are more highly expressed in decayed wood extract (DW) compared to acetate (AC). Negative values represent genes that are down-regulated in DW versus AC.
<sup>b</sup> Homologs in *R. palustris* TIE-1 are upregulated in response to copper [65].

We also observed that several genes involved in metal transport and homeostasis had significant fitness defects in medium containing decayed wood extract and had no fitness defects in medium containing acetate as a sole carbon source (Figure 3, Table 4). Among these, a putative zinc/manganese transporter (RPA0858-RPA0860), the copper efflux transporter *cusA*, and RPA1660, which encodes a copper P-type ATPase, were required for growth in late decay extract (Figure 3 and Table 4). Additionally, transposon insertions in *lgt*, which encodes a prolipoprotein diacylglyceryl transferase, were also identified as having a fitness defect in decayed wood extract (Figure 3 and Table 4), and *lgt* mutants in *Salmonella typhimurium* and *Escherichia coli* are more sensitive to copper [44].

**Table 4.** Fitness scores for selected *R. palustris* genes from RB-Tnseq experiments in minimal medium with acetate compared to minimal medium with decayed wood extract.

| RPA# (locus tag) | gene name | Annotation | DW<sup>a</sup> | Ac<sup>b</sup> |
| --- | --- | --- | --- | --- |
| *Fitness neutral in acetate and fitness loss (or gain) in decayed wood extract* |  |  |  |  |
| RPA0858<br>(TX73_004425) | *aztA* | Zinc ABC transporter ATP-binding protein | -1.57 | -0.74 |
| RPA0859<br>(TX73_019145) |  | Metal ABC transporter permease | -2.06 | -0.84 |
| RPA0860<br>(TX73_004435) |  | Metal ABC transporter substrate-binding protein | -2.55 | -0.6 |
| RPA0980<br>(TX73_005050) | *hoxJ* | PAS domain S-box protein | 6.5 | 0.05 |
| RPA1420<br>(TX73_007285) | *cusA* | Cu-specific metal efflux RND exporter | -1.01 | -0.11 |
| RPA1660<br>(TX73_008535) |  | Heavy metal translocating P-type ATPase | -1.8 | -0.02 |
| *Fitness loss in acetate and fitness neutral in decayed wood extract* |  |  |  |  |
| RPA0254<br>(TX73_001320) | *gpsA* | NAD(P)-dependent glycerol-3-phosphate dehydrogenase | 0.28 | -3.58 |
| RPA0443<br>(TX73_002300) |  | helix-turn-helix domain-containing protein | -0.52 | -1.28 |
| RPA1076<br>(TX73_005530) |  | FAD-binding oxidoreductase | 0.35 | -2.8 |
| RPA1559<br>(TX73_008010) | *cbbL* | form I ribulose biphosphate carboxylase large subunit | 0.39 | -2.14 |
| RPA1560<br>(TX73_008015) | *cbbS* | ribulose biphosphate carboxylase small subunit | 0.48 | -1.4 |
| RPA1561<br>(TX73_008020) | *cbbX* | CbbX protein | 0.56 | -1.58 |
| RPA2045<br>(TX73_010535) | *bioB* | biotin synthase BioB | 0.23 | -1.44 |
| RPA2555<br>(TX73_013215) |  | glycosyltransferase family 39 protein | -0.92 | -1.43 |
| RPA2694<br>(TX73_013960) | *pdxJ* | pyridoxine 5'-phosphate synthase | 0.13 | -2.89 |
| RPA2970<br>(TX73_015405) | *bioA* | adenosylmethionine--8-amino-7-oxononanoate transaminase | 0.34 | -1.4 |
| RPA2971<br>(TX73_015410) | *bioD* | dethiobiotin synthase | 0.19 | -1.98 |
| RPA3048<br>(TX73_015770) | *ppk* | RNA degradosome polyphosphate kinase | 0.64 | -2.1 |
| RPA3065<br>(TX73_015855) | *pdxA* | 4-hydroxythreonine-4-phosphate dehydrogenase PdxA | -0.42 | -2.35 |
| RPA3702<br>(TX73_019180) | *methH* | methionine synthase | 0.75 | -2.14 |
| RPA4661<br>(TX73_024230) | *otsB* | trehalose-phosphatase | -0.28 | -2.03 |
| RPA4748<br>(TX73_024670) | *hbdA* | 3-hydroxybutyryl-CoA dehydrogenase | 0.2 | -1.38 |

<sup>a</sup> fitness scores for genes from the barcoded transposon library grown in photosynthetic medium
(PM) supplied with the decayed wood extract (decayed extract, DE) as a carbon source.
<sup>b</sup> fitness scores for genes from the barcoded transposon library grown in photosynthetic medium
(PM) supplied with 20 mM acetate (Ac) as a carbon source.

To examine the concentration of various metals present in the early and late decayed wood extract which could be responsible for the observed metal stress in *R. palustris*, we used inductively-coupled plasma – mass spectrometry (ICP-MS). Because cation concentrations will increase due to loss of wood mass (denominator), rather than gain of cations (numerator), it is important to adjust mass-based analyses to compensate for this, known as the ‘illusion of gain’ [45]. Regarding the mass adjusted cation data, late decay wood extract had far more Al, Fe, and Cu than the undecayed wood, indicating significant import by the fungus (Figure 4). While *R. palustris* can tolerate concentrations of Fe much higher than this while performing iron oxidation [46] the concentrations of Al and Cu are significantly higher than what is normally found in our standard growth medium [47]. The importance of the metal transport genes, along with evidence that aluminum and copper are relatively concentrated by *R. placenta*, indicates that tolerating elevated metal concentrations is critical for bacterial growth in wood decayed by *R. placenta*.

**Figure 4.**
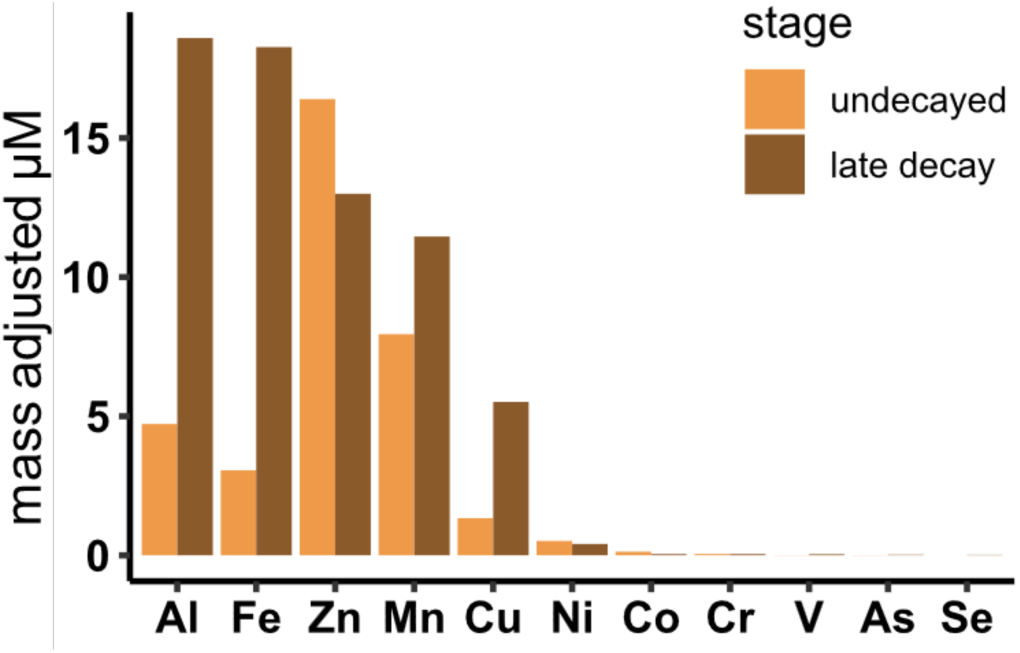
ICP-MS analysis of metal concentrations in undecayed (orange) versus late decay (brown) wood extract. 500 μL of undecayed or late decay wood extract was used for metal identification. Values represent 40 total scans of each sample (see methods) and are adjusted to compensate for wood density loss by multiplying absolute values by the fraction of mass remaining in decayed samples, relative to original.

## Discussion

The purpose of this study was to use the genetically tractable bacterium *R. palustris* to determine how and to what extent an Alphaproteobacterium could thrive on fungal-degraded wood as a carbon source. We found that wood decayed by a model brown rot fungus*, R. placenta,* generates carbon substrates and other metabolites that support bacterial growth, but that lignin residues were not a primary resource for *R. palustris*. These findings support the idea that the activity of wood-degrading fungi can create nutrient niches that could sustain bacterial populations within decaying wood. At the same time, we found that wood decay by *R. placenta* increases the local abundance of certain metals, including toxic metals such as aluminum and copper, indicating that managing metal toxicity could be a critical factor for bacterial survival in wood decay environments, depending on how metal ions are presented (e.g., oxidation state, complexation). Together, these findings demonstrate how a model bacterium can be used to reveal both beneficial and antagonistic aspects of fungal–bacterial interactions in systems that remain largely inaccessible to direct genetic interrogation.

*R. palustris* has attracted biotechnological interest for its ability to degrade aromatic compounds derived from chemically hydrolyzed lignocellulose, and our analyses revealed the presence of carbon substrates that *R. palustris* can assimilate in the decayed wood extract, including aromatic compounds [13,33]. However, aromatic compound degradation does not appear to play a significant role in the growth of *R. palustris* in naturally decayed extracts. The main pathway for aromatic compound degradation in *R. palustris* was not required for growth with fungal-decayed wood extract. Most of the aromatic compounds detected in the late decay extracts were still detected after 95 days of incubation with *R. palustris*. Additionally, most of the genes involved in aromatic compound degradation in *R. palustris* were not induced during growth with decayed wood extract, despite the presence of aromatic compounds known to induce their expression.

This contrasts with previous observations that *R. palustris* can co-utilize acetate and aromatics in corn stover hydrolysate [13]. One potential explanation could be that the lignin composition of corn stover and aspen (being a monocot and a dicot, respectively) is different, where corn has higher p-hydroxyphenyl (H-type) lignin content while aspen is dominated by G- and S-type lignin. However, monocots also contain G-, and S-type lignin in addition to H-type, so the presence of G- and S-type lignin alone does not explain the difference in bacterial metabolism. Another explanation could relate to the fact that aromatic compound degradation in *R. palustris* is strongly inhibited by the presence of long-chain fatty acids, such as linoleic acid [48]. Linoleic acid and related fatty acids are abundant components of fungal phospholipids, and phospholipid fatty acids have often been used as biomarkers to assess fungi, including *R. placenta* [49]. These phospholipid fatty acids may act as inhibitors of bacterial aromatic compound degradation in this system.

In contrast, short-chain organic acids appear to play a central role in bacterial growth in decayed wood extracts. *R. palustris* is known for its ability to co-utilize short-chain organic acids [50], and our data indicate that *R. palustris* can use multiple short-chain organic acids for growth in decayed wood extracts. Acetate, for example, can constitute up to 5% of wood mass in dicot tree species as acetyl ester linkages with hemicellulose and cellulose [51]. Acetate is significantly more abundant in the late decay stage of brown rot, including decay by *R. placenta*, and was detected in late-stage decay extracts in our experiment but was depleted following bacterial growth, consistent with its assimilation by *R. palustris* [12,36]. Brown rot fungi are also known for their ability to secrete oxalate to help chelate Fe^3+^ during the initial stages of decay, which could be a growth substrate for a bacterial partner [52]. Although oxalate was not detected in our extracts, the absence of oxalate could be due to the activity of oxalate decarboxylase, an oxalate-degrading enzyme upregulated in *R. placenta* during later decay stages [53], or could be bound as insoluble calcium oxalate crystals which would have been filtered out during our extraction procedure [54]. Oxalate decarboxylase converts oxalate to formate and CO_2_, and formate is more abundant in the late stage of wood decay by *R. placenta* [12]. However, *R. palustris* is unable to use formate as a sole carbon source, and whether it can be co-utilized with other substrates has not been determined [55]. Instead, we detected oxaloacetate, a precursor to oxalate [52] which was depleted following bacterial growth too, suggesting that it may serve as an additional growth substrate.

Beyond carbon metabolism, RB-TnSeq analyses identified several genes that were dispensable for growth in decayed wood extract but essential in minimal medium with acetate as the sole carbon source. These included genes involved in the biosynthesis of the vitamins biotin and pyridoxal 5′-phosphate, the amino acid methionine, and the metabolites trehalose and glycerol-3-phosphate. This indicates that *R. palustris* can acquire these metabolites from the decayed wood extract, implicating them as potential mediators of fungal-bacterial interactions within wood decay communities. Consistent with this interpretation, trehalose and glycerol-3-phosphate have been detected during late-stage wood decay by *R. placenta*, with glycerol-3-phosphate being significantly more abundant at later stages than in early decay stages [12].

While brown rot activity clearly releases nutrients that can support bacterial growth, wood decay fungi are also thought to possess strategies to suppress microbial competitors. While there is some indication that antagonistic interactions occur between wood rot fungi and bacteria, potential inhibitors have yet to be identified [56,57]. In this context, our observation that *R. palustris* is responding to elevated metal content in decay extracts is noteworthy. Wood decay by *R. placenta* has been reported to lead to a greater abundance of aluminum, iron, and copper in decayed wood [45,52,53,58]. Consistent with elevated metal concentrations causing bacterial stress, we found genes encoding copper efflux systems, including *cusA* and RPA1660, were essential in *R. palustris*, indicating that copper homeostasis is important for bacterial survival in this environment. The antimicrobial properties of copper are well established, and copper can be used as wood preservative and fungicide [59–61]. Because several brown rot fungal species, including *R. placenta*, exhibit a high tolerance for copper [54], likely by trapping and ‘deactivating’ it as insoluble copper oxalate, their ability to increase copper concentrations could serve as a strategy to inhibit bacterial or fungal competitors.

Finally, we observed induction of the GTA system in *R. palustris* during growth with late-decay stage wood extract. GTAs are typically induced under conditions of nutrient limitation and are regulated by quorum sensing, leading to the lysis of a subpopulation of cells and the release of phage-like particles that interact with closely related bacteria as their recipient [62,63]. GTAs have been implicated in DNA repair and genetic exchange, and they are particularly prevalent among Alphaproteobacteria, a group commonly associated with brown rot fungi [2,43]. In a survey of 1,423 Alphaproteobacteria genomes, GTAs were found in almost 60% of the genomes [64]. The induction of the GTA system in this context suggests that wood decay environments may promote horizontal gene transfer or stress-associated genetic exchange, potentially influencing bacterial adaptation and community dynamics during late stages of fungal decay.

Most importantly, this work demonstrates the utility of pairing a genetically-tractable bacterium with high-throughput functional genomics to interrogate microbial interactions in systems that are otherwise experimentally intractable. Our findings raise new questions about how fungal-mediated nutrient release, metal mobilization, and stress-induced genetic exchange shape bacterial evolution and community assembly during wood decay. It also begs the question of whether other bacteria, including other genetically-tractable proteobacteria like *Pseudomonas putida*, or tractable models of other taxa found in decay communities, might be similarly tested to explore new dimensions of fungal-bacterial interactions through the process of fungal wood decay. In summary, we believe that this study can provide a framework for better understanding the fungal-bacterial interactions in the decayed wood environment.

## Acknowledgements

The authors would like to thank Dr. Anthony Zmuda for assistance with operation of the LC-MS and interpretation of data, and Molly Moran and Sarah Anglin for assistance with the wood decomposition microcosms and acid hydrolysis experiments. Element analysis (ICP-MS) was performed at the Northwestern University Quantitative Bulk-Element Information Core (RRID:SCR_017773).

## Study funding

Funding to JSS, KRF, and IUH provided by the University of Minnesota College of Biological Sciences (CBS) and through a Biocatalysis Initiative award by the BioTechnology Institute, University of Minnesota. Support for NML provided by award DE-SC0020252 from the U.S. Department of Energy, Office of Science, Basic Energy Sciences, Physical Biosciences program awarded to KRF.

## CRediT roles

Nathan Lewis: Data curation, formal analysis, investigation, methodology, visualization, writing the original draft, reviewing and editing the manuscript

Irshad Ul Haq: Data curation, conceptualization, investigation, methodology, resources, visualization, reviewing and editing the manuscript

Jonathan Schilling: Conceptualization, resources, supervision, reviewing and editing the manuscript

Kathryn Fixen: Conceptualization, project administration, resources, supervision, validation, reviewing and editing the manuscript

## Notes

### Competing Interest Statement

The authors have declared no competing interest.

