## Supplemental methods for "Leveraging a genetically tractable alphaproteobacterium reveals molecular determinants of bacterial growth in fungal-decayed wood"

*Cultivation of the barcoded transposon library*

Aliquots of the barcoded transposon library used in [1] were thawed from -80°C on ice and inoculated into 50 mL of PM supplemented with 0.2% yeast extract and 0.5% casamino acids with 20 mM acetate as a carbon source and 100 mg/mL kanamycin to maintain selection for the transposable element. The library was grown phototrophically in anoxic conditions until the culture reached mid logarithmic growth phase. 1 mL of the library in mid-log phase was removed, and cells were harvested by centrifugation. The supernatant was discarded and the cell pellet was frozen at -80 °C and was used to define the initial barcode abundance in the library.

The remainder of the library was used to inoculate 10 mL liquid cultures of anoxic PM with decayed wood extract as a carbon source and PM with 20 mM acetate as a control, both in biological triplicate as follows. 1 mL of the library was removed from the 50 mL culture and cells were pelleted by centrifugation. The supernatant was discarded and the cells were washed twice with PM lacking any carbon source, then resuspended in PM to an $OD_{660}$ of 1.0. The washed cells were then used to inoculate 10 mL cultures of PM with acetate or PM with decayed wood extract in biological triplicate to a starting $OD_{660}$ of 0.03 in septum-sealed anaerobic culture tubes. Cultures were incubated in the light as described above until they reached an optical density at 660 nm of 0.6. Afterwards, 1 mL of each culture was harvested by centrifugation. The supernatant was discarded and the cell pellet was frozen in liquid nitrogen. Samples were shipped to Lawrence Berkely National Laboratory for lysis, sequencing, and barcoded transposon quantification as described in [1]. The barcode abundance for each gene in PM with acetate and PM with late decay wood extract is available as supplemental dataset 2 (Table S1) and can be viewed using the Fitness Browser (https://fit.genomics.lbl.gov/cgi-bin/org.cgi?orgId=RPal_CGA009).

*RNA extraction and sequencing*

RNA was extracted from cultures of *R. palustris* grown in anoxic PM with acetate and PM with decayed wood extract as carbon sources in biological triplicate. The *R. palustris* cultures grown using decayed wood extract used for RNA extraction and sequencing are those shown in Figure 2. RNA was harvested after 95 days of phototrophic growth. *R. palustris* cultures grown on PM with acetate were also grown to early stationary phase and had an $OD_{660}$ between 0.9 and 0.99 at the time they were harvested. For both conditions, cells were harvested by centrifugation, then the supernatant was removed and the cell pellets were flash-frozen in liquid nitrogen, and stored at -80 °C. The supernatant of cultures grown in PM was kept for LC-MS analysis described later.

RNA was purified from liquid cultures of *R. palustris* using the miRNeasy Mini Kit (Qiagen) to purify total RNA, and the RNeasy MinElute cleanup kit to purify DNA-depleted RNA with minor modifications to the manufacturers procedures as described previously [2,3]. The resulting RNA was shipped to SeqCenter (Pittsburgh, PA) for library preparation and sequencing. For cultures grown in PM with acetate and PM with decayed wood extract, two biological replicates were sequenced.

cDNA library synthesis was performed using the Illumina stranded Total RNA prep ligation kit and ribosomal RNA was removed using the Illumina Ribo-Zero plus kit. 10 bp unique dual indices were added to the resulting DNA fragments. Sequencing of the cDNA library was performed using NovaSeq X Plus to generate 150 bp paired end reads. Demultiplexing, quality control, and adapter trimming was performed with bcl-convert version 4.2.4 (Illumina). Reads were mapped to the published genome of *R. palustris* CGA009 (NCBI: NZ_CP116810) using HISAT2 version 2.2.0 [4] using the 'very sensitive' mode. Reads were quantified using Subread's featureCounts function, version 2.0.1 [5] using the default parameters. Read counts were loaded into Rstudio (version 4.3) and differential gene expression analysis was performed using DEseq2 (version 1.42.1) [6] with a minimum read cutoff of 10 reads in both sequenced replicates and a statistical significance threshold of 0.05. *p*-values were corrected for multiple comparisons using the Benjamini-Hochberg method to provide the adjusted *p*-value presented in supplemental data (Table S2).

*Wood mass loss and acid hydrolysis*

Wood mass loss, a measure of decay progress that can be qualified as a "decay class" (scale 1-5) [7,8] for individual tree species, was measured by comparing the oven-dried (100 °C for 48 hours) mass of wood wafers decayed by *R. placenta* with undecayed wafers as a control. The enrichment of acid-insoluble (i.e., Klason) lignin in late decay *R. placenta*-decomposed wafers was determined using the following procedure: Oven dried wafers were first milled to 20 mesh using a Wiley mill. 60 mg of milled wood was then transferred to 25 mL Balch tubes and 0.6 mL of 72% sulfuric acid was added. The contents of the tubes were thoroughly mixed with glass stir rods until biomass was saturated. The tubes were incubated at 30˚C in a water bath for 60 minutes and mixed with a glass stir rod every 5 to 10 minutes. Afterwards, 16.8 mL of deionized water was added to each tube and tubes were sealed with rubber septa and crimped with an aluminum seal, then autoclaved for 1 hour at 121˚C and 103 kPa. The autoclaved acid hydrolysis solutions were vacuum-filtered using Büchner funnels equipped with glass fiber filters (Fisherbrand Glass Fiber Circles - G6) which had previously been oven-dried (100 ˚C for 2 hours) and weighed. The filtered solids (acid-insoluble) were oven-dried for 24 hours at 100˚C and the mass was measured using a digital scale.

*Identification of compounds present in decayed wood extracts*

Liquid chromatography – mass spectrometry (LC-MS) was used to identify compounds present in the undecayed, early decay, and late decay wood extracts. Standards of lignin monomers and non-aromatic acids suspected to be present in the wood extract were prepared to help identify signals from the very diverse wood extract LC-MS profile. Standards of sodium acetate, sodium fumarate, sodium benzoate, sodium 4-hydoxybenzoate, sodium coumarate, vanillic acid, vanillin aldehyde, cinnamic acid, and sinapic acid were made at a concentration of 50 mM. All samples and standards used for LC-MS were dissolved in HPLC grade $H_2O$ (Millipore Sigma) except for sinapic acid and vanillin, which were dissolved in 100% ethanol. All standards were then diluted 1:100 into HPLC grade water and transferred to HPLC vials prior to being loaded onto the column.

Samples from culture supernatant were prepared by collecting 500 µL of culture supernatant after 95 days of incubation. Supernatant was transferred to a 3 kDa molecular weight cutoff centrifugal filter unit (Millipore Sigma) to remove any cells, cell debris, contaminating proteins, or larger lignocellulose fragments. Flowthrough from the centrifugal filter units was stored at 4 °C until they could be analyzed using LC-MS (<1 month).

All LC-MS analyses were performed on an Agilent 1260 Infinity II and single quadruple MSP equipped with a 2.1 x 50 mm reverse-phase SB-C18 column (1.8 µm). 5 µL of each sample or standard were loaded onto the column using an autoinjector which was rinsed with 100% isopropyl alcohol between each load. 0.1% formic acid was used as the mobile phase for liquid chromatography at a flow rate of 0.4 mL $min^{-1}$. Eluate was ionized by electrospray ionization and mass spectrometry was run in tandem positive and negative mode with a capillary voltage of 3000 V. LC-MS enabled the detection of multiple *m/z* peaks for each standard which generally corresponded to the $[M +/-H]^{+/-}$, $[M*Na]^{+/-}$, and $[2M*Na]^{+/-}$ signals although other assignments were occasionally used. The mass spectra for each sample and standard can be found on Figshare under the DOI 10.6084/m9.figshare.32002746.

*Metal identification and quantitation in decayed wood extracts*

Undecayed and late decay wood extracts were submitted to the Quantitative bulk-elemental information core (QBIC) at Northwestern University for metal quantitation analysis. Quantification of Al, V, Cr, Mn, Fe, Co, Ni, Cu, Zn, As, Se in undecayed and late decay wood extracts was accomplished using Inductively-Coupled Plasma – Mass Spectrometry (ICP-MS). Specifically, 0.5 mL of wood extract was heated with 0.5 mL nitric acid ($HNO_3$, > 69%, Thermo Fisher Scientific, Waltham, MA, USA) at 65 °C for 4 hours until completely digested. The resulting solution was diluted with 9 mL ultra-pure water (18.2 MΩ·cm). Quantitative standards were prepared by mixing qualified single-element standards from Inorganic Ventures (Christiansburg, VA, USA) in 5% nitric acid (v/v) to create a calibration curve in a range appropriate for each element.

ICP-MS was performed on a computer-controlled (QTEGRA software) Thermo iCapQ ICP-MS (Thermo Fisher Scientific, Waltham, MA, USA) operating in KED mode and equipped with an electro-spray ionization SC-2DX PrepFAST autosampler (Omaha, NE, USA). Nickel skimmer and sample cones were used from Thermo Scientific (part numbers 1311870 and 3600812). Internal standard was added inline using the prepFAST system and consisted of 1 ng $mL^{-1}$ of a mixed element solution containing Bi, In, $^{6}Li$, Sc, Tb, Y (IV-ICPMS-71D from Inorganic Ventures). Each sample was acquired using 1 survey run (10 sweeps) and 3 main runs (peak jumping) for 40 sweeps total. The isotopes selected for analysis were $^{27}Al$, $^{51}V$, $^{52}Cr$, $^{55}Mn$, $^{56,57}Fe$, $^{59}Co$, $^{60}Ni$, $^{63,65}Cu$, $^{64,66,68}Zn$, $^{75}As$, $^{77}Se$ with $^{45}Sc$, $^{89}Y$, $^{115}In$ included as internal standards for data interpolation and machine stability.
